# Peripheral Immune Cell Shifts Associated With Alzheimer’s Disease Vary in Multi-Ancestry Cohorts

**DOI:** 10.64898/2026.11.11.765813

**Authors:** Yousef Mustafa, Yuchen Yang, Makaela Mews, Tianjie Gu, Lissette Gomez, Larry D. Adams, Takiyah D. Starks, Concepcion Silva, Katalina F. McInerney, Jeffery M. Vance, Michael L. Cuccaro, Maryenela Illanes-Manrique, Mario R Cornejo-Olivas, Briseida E. Feliciano-Astacio, Goldie S. Byrd, Margaret A. Pericak-Vance, Jonathan L. Haines, Anthony J. Griswold, William S. Bush

**Author notes:** **Corresponding Authors**: William S. Bush, Yousef Mustafa.

## Abstract

**Background:** Alzheimer’s disease (AD) is heavily influenced by immune system dysregulation, yet the role of peripheral immune processes—and how these vary across ethnicities—remains understudied.

**Objective:** To investigate ancestry-specific immune alterations in AD by profiling peripheral immune cell proportions using high-resolution deconvolution of whole blood RNA-sequencing data, characterizing disease associated cell types, and identifying enriched biological pathways.

**Methods:** We analyzed whole blood RNA-seq data from non-Hispanic White (NHW), African American (AA), Caribbean Hispanic (CariHISP), and Peruvian (PERU) individuals, balanced between AD cases and cognitively normal controls. Immune cell proportions were estimated using CIBERSORTx and compared using linear models (*propeller* R package) adjusted for age, sex, and APOE genotypes in case-control models. Cell-type-specific differential expression was performed using CARseq, and pathway enrichment analyses performed using MSigDB gene sets.

**Results:** Baseline immune composition differed substantially across cohorts for multiple innate and adaptive cell types. In a multi-ethnic AD versus control analysis, eight cell types reached FDR significance: AD was associated with decreased proportions naïve B cells, CD56dim NK cells, B cells, intermediate monocyte I, and double-negative T cells, alongside significant expansion in neutrophils and non-classical monocytes. Interaction analyses revealed heterogeneous group-specific effects, with the broadest shifts in AA and CariHISP. Pathway enrichment indicated shared heme metabolism, interferon, and oxidative phosphorylation programs across groups, with distinct emphases: broad, strong heme metabolism enrichment and the highest oxidative phosphorylation signal (in myeloid populations) in AA; MYC targets in naïve B cells and eosinophils in CariHISP; dendritic cell–and T follicular helper cell–centered interferon, inflammatory, and proliferative programs in NHW; and TGF-β and apoptosis signatures, unique to PERU, alongside the broadest cell-type coverage.

**Conclusions:** Peripheral immune dysregulation in AD is characterized by depletion of lymphoid and intermediate monocyte populations with concurrent neutrophil and non-classical monocyte expansion, and these patterns vary meaningfully by ancestry. These findings underscore the need for population-informed biomarkers and therapeutic strategies in AD.

## Introduction

Alzheimer’s disease (AD) is a progressive neurodegenerative disorder and the leading cause of dementia worldwide.^1,2^ Its classical neuropathological features include extracellular amyloid-beta (Aβ) plaques and intracellular neurofibrillary tangles composed of hyperphosphorylated tau protein (pTau), first described by Aloïs Alzheimer in 1907.^3^ For decades, the amyloid cascade hypothesis dominated the field, proposing that Aβ accumulation initiates a sequence of events—including microglial activation, tau phosphorylation, synaptic dysfunction, and ultimately, neurodegeneration.^4,5^ However, numerous failed clinical trials targeting Aβ clearance, and observations of cognitively normal individuals with substantial Aβ burden, have challenged this model.^5–9^ These findings suggest that Aβ deposition alone is not sufficient to cause the disease and that other biological processes, particularly inflammation, may play a more central role.

Neuroinflammation is now recognized as a major driver of AD progression.^10–12^ Microglia, the brain’s resident immune cells, are activated by Aβ through pattern recognition receptors such as TLRs and RAGE, leading to the release of pro-inflammatory cytokines, oxidative stress, and further neuronal injury.^13^ While initially protective, chronic microglial activation becomes maladaptive, contributing to sustained inflammation and impaired clearance of toxic proteins. This inflammatory response may be amplified by dysfunction in astrocytes and oligodendrocytes, creating a toxic environment that promotes tau pathology and synaptic loss.^14^ Experimental evidence suggests that neuroinflammation not only accompanies but may precede Aβ accumulation and tau hyperphosphorylation, broadening the conceptual framework to include inflammation-driven disease pathways.^13,15^

Compelling data now support a systemic dimension to this neuroinflammation.^16–18^ Blood-brain barrier (BBB) disruption in AD facilitates the migration of peripheral immune cells—including monocytes, neutrophils, and T cells—into the brain, where they can contribute to neuroinflammatory cascades.^19^ Simultaneously, inflammatory mediators produced in the brain, such as IL-1β and TNFα, are released into the circulation, where they activate peripheral immune cells and propagate a feedback loop of immune activation.^17,19,20^ The choroid plexus and meningeal lymphatics have emerged as key anatomical gateways for this peripheral–central interaction, mediating immune cell trafficking and modulating CNS immune tone.^20,21^ In this context, AD should be viewed not solely as a brain disorder, but as a condition with systemic immune involvement, wherein peripheral innate immunity plays a critical role in disease onset and progression.

Among peripheral immune cells, monocytes and neutrophils have received increasing attention.^22–28^ Monocytes are capable of phagocytosing Aβ and presenting antigens via MHC class II, yet their function appears impaired in AD, with reduced phagocytic capacity and altered expression of TLRs and chemokine receptors.^22^ Neutrophils, although short-lived, have been implicated in early-stage pathology and can exacerbate tissue damage through excessive production of reactive oxygen species.^26^ In addition, adaptive immune cells such as T– and B-lymphocytes have shown substantial effects on AD progression through their influence on microglial activation, even from the periphery.^29,30^ Despite these findings, the systemic immune alterations in AD remain incompletely characterized, particularly in relation to central-peripheral immune interactions and how they may be leveraged for novel treatment approaches.

Furthermore, ancestry significantly influences immune system architecture and responsiveness.^31^ Differences in immune cell proportions, receptor polymorphisms, and cytokine signaling pathways across populations may alter susceptibility to neurodegeneration and the trajectory of AD-related immune responses.^32^ Yet, most transcriptomic and immunological studies in AD have been conducted in cohorts lacking ethnic diversity, limiting their relevance for global populations. This lack of inclusion also hampers the development of precision medicine approaches that consider population-specific risk profiles and therapeutic responses.

This study addresses these critical gaps by profiling peripheral immune cell composition in AD using high-resolution deconvolution of whole blood RNA-sequencing data from a multi-ancestry cohort including non-Hispanic White (NHW), African-American (AA), Peruvian (PERU), and Caribbean Hispanic (CariHISP) individuals. We aim to identify immune cell types associated with AD and evaluate how these associations differ by ancestry. By uncovering both shared and population-specific immune alterations, our findings seek to inform the design of population-specific biomarkers and therapeutic strategies for AD.

## Materials and Methods

### Study Population

This study used data from the Multi-Ancestry Genomics, Epigenomics, and Transcriptomics of Alzheimer’s (MAGENTA) Project,^33^ a component of the Alzheimer’s Disease Sequencing Project (ADSP).^34^ A total of 919 individuals were enrolled across three institutions: the John P. Hussman Institute for Human Genomics at the University of Miami Miller School of Medicine (Miami, FL), North Carolina A&T State University (Greensboro, NC), and Case Western Reserve University (Cleveland, OH), as part of the ADSP Follow-up Study. Participants included both Alzheimer’s disease (AD) cases and cognitively normal controls, all aged 65 years or older at the time of disease onset (for cases) or clinical examination (for controls).

Clinical diagnoses of AD were established by expert adjudication panels at each site, using standardized diagnostic criteria developed by the National Institute on Aging and the Alzheimer’s Association. Whole blood was collected from each participant, and RNA was extracted and subjected to high-throughput RNA sequencing. For the present analysis, individuals with mild cognitive impairment (MCI) or other neurological or psychiatric conditions were excluded, leaving only AD cases and cognitively normal controls. After quality control and filtering, the final analytic cohort included 240 non-Hispanic White (NHW) individuals (121 AD cases, 119 controls), 234 African-American (AA) individuals (115 AD cases, 119 controls), 294 Caribbean Hispanic (CariHISP) individuals (143 AD cases, 151 controls), and 88 Peruvian (PERU) individuals (41 AD cases, 47 controls), for a total of 856 samples across all cohorts.

### Cell Type Proportion Estimation And Modeling

To investigate population-specific immune alterations associated with Alzheimer’s disease (AD), we estimated immune cell type proportions from whole blood RNA-sequencing (RNA-seq) data using CIBERSORTx,^35^ a high-resolution digital cytometry tool. The analysis was conducted across four ethnic groups: non-Hispanic White (NHW), African-American (AA), Caribbean Hispanic (CariHISP), and Peruvian (PERU) participants.

Cell type deconvolution was performed using the 1M-scBloodNL single-cell RNA-seq reference panel,^36^ developed by the eQTLGen consortium, as well as the LM22 panel developed by CIBERSORT. The single-cell reference matrix includes gene expression profiles derived from peripheral blood mononuclear cells (PBMCs) of European individuals and enables high-resolution identification of diverse immune cell types, including subsets of monocytes, B cells, T cells, and natural killer (NK) cells. The 1M-scBloodNL reference is particularly well-suited for resolving myeloid lineages and allowed us to quantify distinct monocyte populations, such as classical, non-classical, and intermediate monocytes. Deconvolution was run independently within each ethnic group and for each reference panel, with adjusted cell fraction estimates merged by sample. Where cell types with the same names was estimated from both reference panels (e.g., naïve CD4□ T cells and regulatory T cells), the panel is indicated with (LM22) or (1M) in the Figures and text.

To assess how immune cell composition differed by AD status and ancestry, we used the *propeller* R package to model logit-transformed cell type proportions using linear models with robust empirical Bayes moderation.^37^ All named cell types from both reference panels, together with the unannotated “unknown” cell population, were tested in every model.

First, baseline models tested differences in cell type proportions between ancestry groups, with ancestry as the predictor and age and sex as covariates; AD status was not included, and all six pairwise contrasts between groups were tested (**Figure 1**).

**Figure 1.**
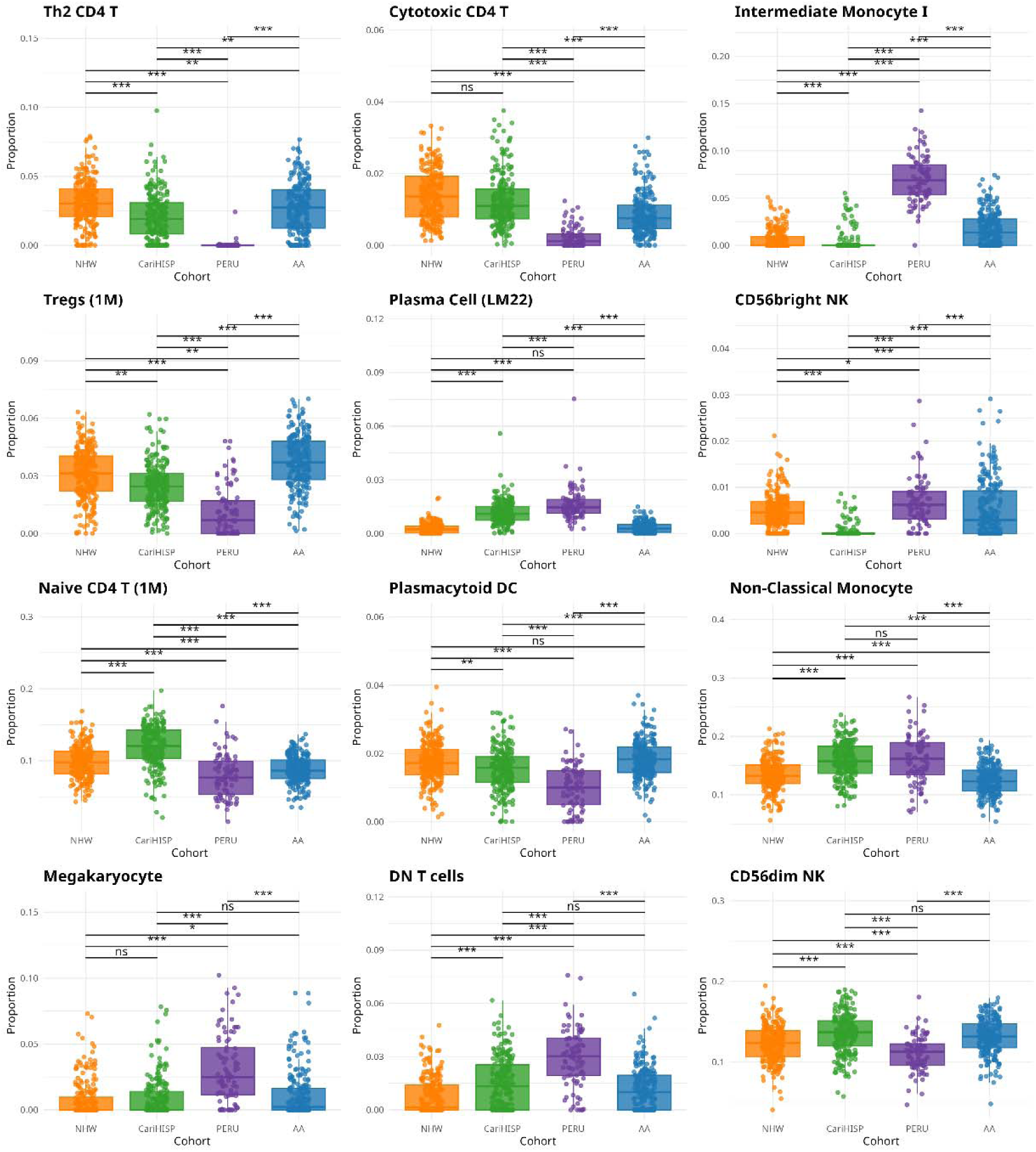
Baseline Peripheral Immune Cell Composition Differs in Multi-Ethnic Cohorts. Baseline peripheral immune cell composition differs across ancestry groups. Boxplots display the estimated proportions of the top 12 representative immune cell types across non-Hispanic White (NHW), African American (AA), Caribbean Hispanic (CariHISP), and Peruvian (PERU) participants at baseline, independent of Alzheimer’s disease status. Cell type proportions were inferred from whole-blood RNA sequencing using CIBERSORTx with the 1M-scBloodNL and LM22 reference panels. Linear modeling was conducted using the *propeller* package, adjusting for age and sex. Black bars denote pairwise comparisons between ancestry groups, with corresponding significance levels shown above each panel. Significance levels are indicated as follows: *p* < *0.05 (*), p < 0.01 (**), p < 0.001 (***)*, and ns = not significant.

Second, a pooled model compared AD cases with controls across all four groups, adjusting for ancestry, age, sex, and APOE e2 and e4 allele counts to estimate the overall association between AD and immune composition (**Figure 2**). Individuals without verified APOE genotypes (n = 1) were excluded from all AD case-control models (n = 855) but retained in the baseline models (n = 856).

**Figure 2:**
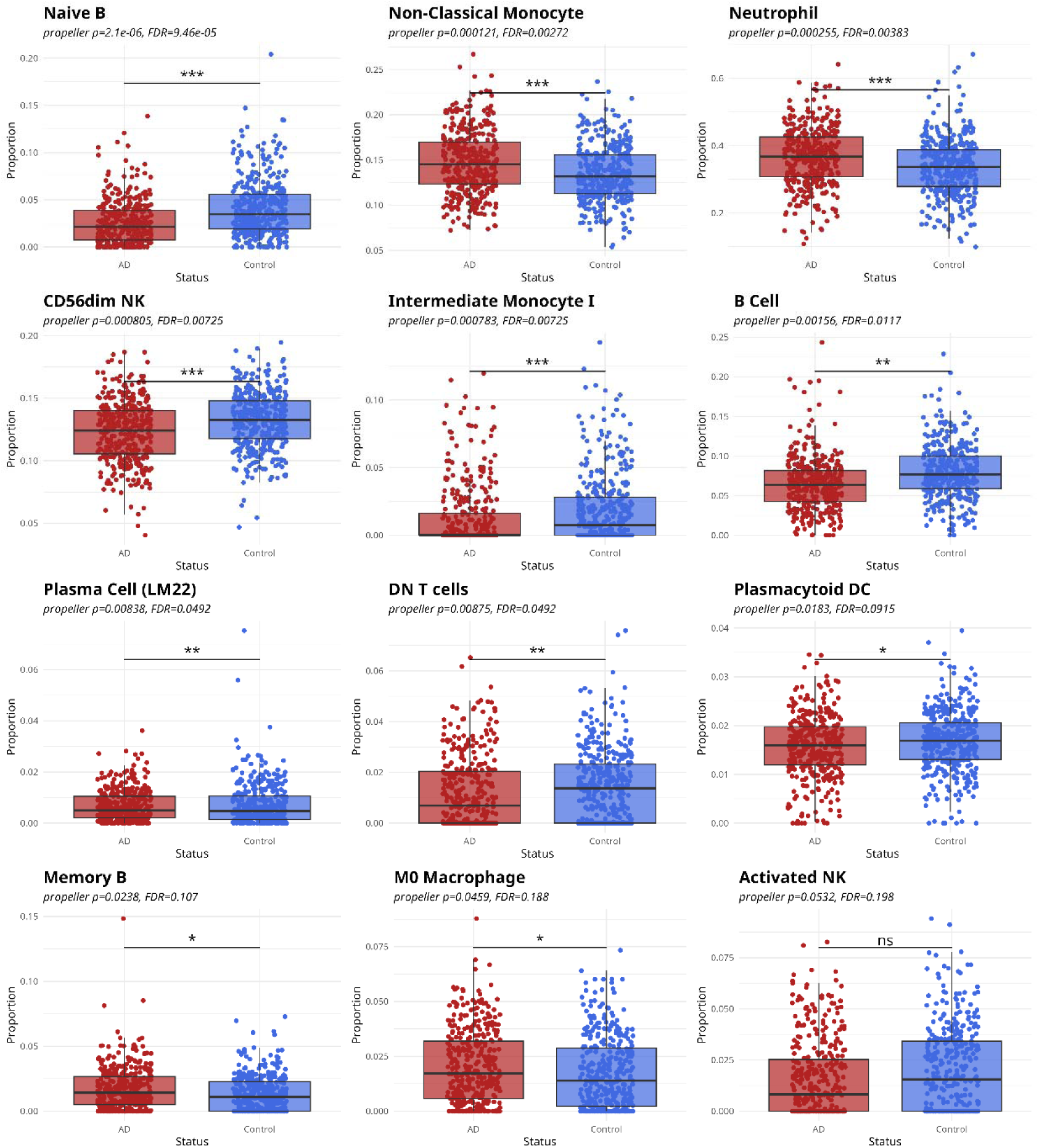
Multi-Ethnic Analysis of Peripheral Immune Cell Proportions Reveals Disease-Associated Shifts in AD. Boxplots show differences in immune cell type proportions imputed from whole blood RNA-seq using CIBERSORTx. Comparisons between AD cases (red) and controls (blue) were performed using the *propeller* R package, adjusting for age, sex, APOE e2 and e4 allele counts, and ancestry. Significance: *P* < 0.05 (\**), < 0.01 (**), < 0.001 (\*\*\**); FDR values shown.

Third, to estimate ancestry-specific AD effects, we fit a single model in which each combination of AD status and ancestry group (eight groups in total) was represented as a separate level, with age, sex, and APOE e2 and e4 allele counts as covariates shared across groups. This parameterization is equivalent to a model containing main effects of AD status and ancestry and a status-by-ancestry interaction term. From this third model, we extracted two types of contrasts. Within-ancestry AD contrasts compared AD cases with controls separately in NHW, AA, CariHISP, and PERU, testing whether AD was associated with a difference in cell type proportion within each group (**Figure 3**). This approach allowed us to extract ancestry-specific AD effects from a single model, with age, sex, and APOE effects shared across groups and residual variance moderated across cell types.

**Figure 3.**
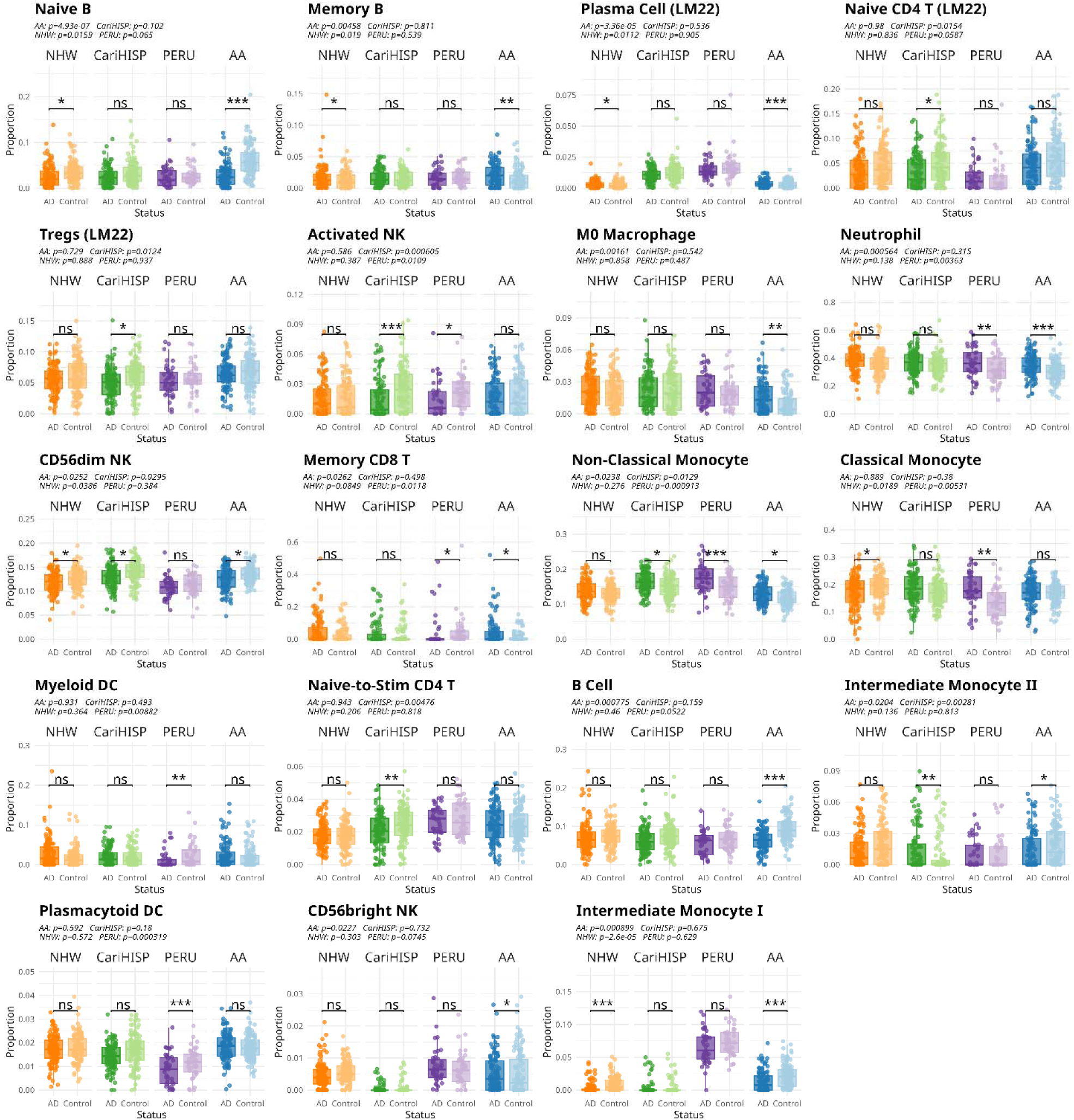
Ethnicity-specific peripheral immune cell shifts associated with Alzheimer’s disease. For each cell type, boxplots compare AD and control participants within each group (NHW, AA, CariHISP, PERU). Points represent individuals; boxes show the interquartile range with median lines. Proportions were deconvolved from whole-blood RNA-seq using CIBERSORTx with the LM22 and 1M-scBloodNL references. Within-group differences (AD vs control) were tested using the *propeller* framework with logit transformation and models adjusted for age, sex, and APOE e2 and e4 allele counts; stars above brackets denote cohort-specific p-values (p < 0.05 (*), < 0.01 (**), < 0.001 (***); ns = not significant).

Interaction contrasts compared the AD-versus-control difference in NHW with that in each of the other three groups ((AD_NHW_ – Control_NHW_) − (AD_group_ – Control_group_)), using NHW as the reference (**Figure 4**). Contrasts among the non-NHW groups were not tested. This approach allowed us to extract ancestry-specific AD effects while leveraging shared variance across the full dataset. Additionally, the interaction term enabled formal testing of whether the impact of AD status on immune composition significantly differed across ancestries. This modeling strategy provided a unified framework to evaluate both shared and ancestry-specific immune alterations in AD.

**Figure 4.**
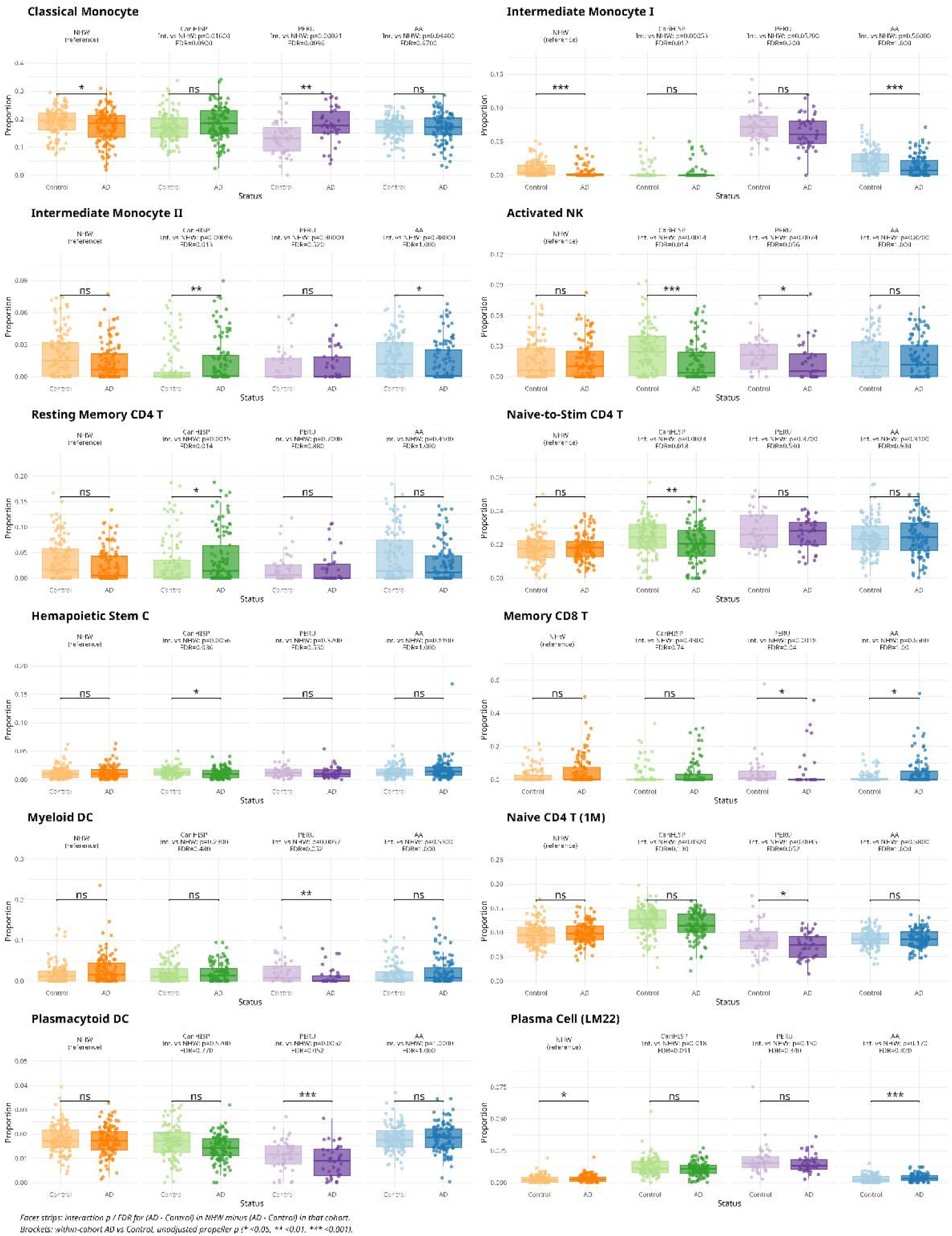
Interaction Model of ancestry and AD Status Reveals Population-Specific Peripheral Immune Alterations in AD. For each cell type, boxplots compare control and AD participants within each group (NHW, CariHISP, PERU, AA). Points represent individuals; boxes show the interquartile range with median lines. Proportions were deconvolved from whole-blood RNA-seq using CIBERSORTx with the LM22 and 1M-scBloodNL references Panel headers report the interaction P-value and FDR for (AD − Control in NHW) − (AD − Control in the indicated group), adjusted for age, sex, and APOE e2 and e4 allele counts; FDR was calculated across all cell types within each contrast. The 12 cell types with the lowest FDR across the three interaction contrasts are shown. Stars above brackets denote unadjusted within-group AD versus control P-values (p < 0.05 (\**), < 0.01 (**), < 0.001 (\*\*\**); ns = not significant).

P-values were adjusted for multiple testing using the Benjamini–Hochberg procedure across all cell types within each contrast, and significance symbols in the Figures reflect unadjusted P-values. Cell types shown in the Figures were selected by FDR: the 12 cell types with the lowest FDR across the six baseline contrasts (**Figure 1**) or in the pooled AD model (**Figure 2**), among those with FDR < 0.20; all cell types with FDR < 0.10 in at least one ancestry-specific AD contrast (**Figure 3**); and the 12 cell types with the lowest FDR across the three interaction contrasts (**Figure 4**). The unannotated “unknown” population was omitted from **Figures 1, 3**, and **4**.

### Cell-Type-Specific Differential Expression Analysis

To identify transcriptional changes within specific peripheral immune cell subsets associated with Alzheimer’s disease (AD), we performed cell-type-specific differential expression (DE) analysis using the CARseq (Cell Type Aware analysis of RNA-seq) framework.^38^ CARseq is a statistical method that models RNA-seq count data with a negative binomial distribution while incorporating cell type proportions to estimate gene expression changes within each cell type independently.

Cell type proportion estimates derived from CIBERSORTx Hi-Res deconvolution, based on the LM22 and 1M-scBloodNL single-cell reference matrices, were used as input to CARseq. These proportions were integrated with raw bulk RNA-seq count data to estimate the contribution of each cell type to the observed gene expression in whole blood. CARseq models were constructed to compare AD cases versus cognitively normal controls, including covariates for age, sex, APOE e2 and e4 allele counts, and surrogate variables to control for batch effects and other latent confounders.

Differential expression analysis was conducted separately for each of the imputed immune cell types across the full cohort as well as within each ethnic group (non-Hispanic White, African-American, Caribbean Hispanic, and Peruvian). Genes were considered differentially expressed at a false discovery rate (FDR) < 0.05. This approach enabled the detection of cell-type-specific gene expression changes associated with AD, accounting for variation in cell type composition and allowing population-specific transcriptional signatures to be resolved. The results were subsequently used for downstream pathway enrichment analysis to elucidate biological processes perturbed in AD across different immune cell types and ethnicities.

### Pathway Enrichment Analysis of Cell-Type-Specific Differential Expression

To investigate the biological processes associated with transcriptional alterations in Alzheimer’s disease (AD), we performed pathway enrichment analysis on differentially expressed genes identified through CARseq across multiple immune cell types and ethnic populations. For each group—non-Hispanic White, African-American, Caribbean Hispanic, and Peruvian—we selected genes with an adjusted p-value below 0.05 and an absolute log2 fold change greater than 1.0. These genes were considered significantly differentially expressed within each cell type.

Enrichment analysis was then conducted using a hypergeometric test framework, implemented via the *clusterProfiler* R package.^39^ Gene set annotations were obtained using the *msigdbr* package, which provided curated collections from the Molecular Signatures Database (MSigDB), including Hallmark pathways (H), Immune Signatures (C7), Canonical Pathways (C2:CP), microRNA Target Sets (C3:MIR), and Human Phenotype Ontology (C5:HPO), as well as external pathway resources such as KEGG and WikiPathways.^40^ Significant pathways were defined using a Benjamini-Hochberg FDR-adjusted p-value < 0.05. For each group, the 15 pathways with the strongest enrichment are shown, with color scaled to −log□□ adjusted P (capped at 10).

By analyzing pathway activity within each ethnic group, this approach allowed us to identify both conserved and divergent biological mechanisms of immune dysregulation in AD, uncovering population-specific patterns of inflammation and immune response that may inform targeted interventions.

## Results

### Baseline Peripheral Immune Cell Composition Varies in Multi-Ethnic Cohorts

We first examined whether circulating immune cell type proportions differed by ancestry, independent of AD disease status, across non-Hispanic White (NHW), African American (AA), Caribbean Hispanic (CariHISP), and Peruvian (PERU) groups.

This analysis revealed widespread ancestry-associated variation in baseline peripheral immune composition. As illustrated in **Figure 1**, the top twelve cell types by FDR demonstrated marked differences between groups, with three broad patterns emerging. First, Th2 CD4□ T cells and cytotoxic CD4□ T cells shared a similar profile, with the highest proportions in NHW, intermediate levels in CariHISP and AA, and notably lower levels in PERU. Second, plasma cells and non-classical monocytes showed an inverse pattern, with the highest proportions in CariHISP and PERU and relative depletion in NHW and AA. Third, regulatory CD4□ T cells, plasmacytoid dendritic cells, intermediate monocyte I, megakaryocytes, and double-negative T cells displayed more nuanced distributions in PERU diverged substantially from all 3 other groups, with markedly elevated intermediate monocytes and megakaryocytes, and reduced plasmacytoid DCs and regulatory CD4□ T cells relative to the other three groups. CD56dim NK cells were most abundant in CariHISP and lowest in PERU. Naïve CD4□ T cells were most abundant in CariHISP, whereas CD56bright NK cells were least abundant in CariHISP.

Overall, these results confirm that peripheral immune architecture varies considerably across ethnic groups, even in the absence of Alzheimer’s disease, and that certain cell types stratify along an NHW/AA versus CariHISP/PERU axis while others highlight the distinctiveness of the Peruvian cohort. These patterns reinforce the importance of population-stratified analyses in immunogenomic studies.

### Multi-Ethnic Analysis of Peripheral Immune Cell Proportions Reveals Disease-Associated Shifts in AD

We next assessed peripheral immune cell shifts associated with AD across all four ethnic groups to identify immune signatures that were consistent and potentially generalizable across populations. By aggregating samples while adjusting for age, sex, APOE e2/e4 allele counts, and ancestry, we aimed to characterize the core peripheral immune features of AD, independent of population-specific effects.

This multi-ethnic analysis revealed broad peripheral immune shifts associated with AD (**Figure 2**). Eight cell types reached FDR significance (FDR < 0.05). The most significant associations were observed for naïve B cells (P = 2.1 × 10□□, FDR = 9.46 × 10□□), non-classical monocytes (P = 1.21 × 10□□, FDR = 2.72 × 10□³), and neutrophils (P = 2.55 × 10□□, FDR = 3.83 × 10□³)—with naïve B cells decreased and non-classical monocytes and neutrophils increased in AD cases relative to controls. CD56dim NK cells (P = 8.05 × 10□^4^, FDR = 7.25 × 10□³), B cells (P = 1.56 × 10□^3^, FDR = 1.17 × 10□²), intermediate monocyte I (P = 7.83 × 10□^4^, FDR = 7.25 × 10□³), and double-negative T-cells (P = 8.75 × 10□^3^, FDR = 4.92 × 10□²) also showed significant reduction in AD. Plasma cells (P = 8.38 × 10□^3^, FDR = 4.92 × 10□²) showed modest elevation in AD after covariate-adjustment, although unadjusted mean proportions were slightly higher in controls, reflecting a small number of controls with high plasma cell proportions.

Additional cell types with nominally altered proportions in AD that did not reach FDR significance included plasmacytoid dendritic cells (P = 0.0183, FDR = 0.0915), memory B cells (P = 0.0238, FDR = 0.107), and macrophages M0 (P = 0.0459, FDR = 0.188). Memory B cells and macrophages M0 were nominally elevated and plasmacytoid dendritic cells nominally reduced in AD.

**Supplemental Table 1** shows the *propeller* results, including raw proportions of each cell type and t-statistic for each comparison after covariate-adjustment.

Collectively, these results suggest that Alzheimer’s disease is characterized by systemic immune remodeling marked by depletion of circulating lymphoid and intermediate monocyte populations, accompanied by neutrophil and non-classical monocyte expansion. The concurrent reductions in both innate (intermediate monocytes) and adaptive (B cell, T cell) compartments, alongside increases in neutrophils and non-classical monocytes, may reflect impaired immune homeostasis, chronic inflammation-driven exhaustion, or redistribution of specific immune populations between peripheral blood and sites of neuroinflammation.

### Ethnicity-Specific Peripheral Immune Cell Shifts Associated with Alzheimer’s Disease

To investigate whether AD-associated changes in peripheral immune composition differ across multi-ethnic cohorts, we developed group-specific *propeller* models of cell-type proportions, adjusted by age, sex, and APOE e2/e4 allele counts (**Figure 3**). This analysis revealed heterogeneity of disease effects within each ethnic group.

In NHW, AD was associated with significant reductions in intermediate monocytes I (*P* = 2.6 × 10□□), naïve B cells (*P* = 0.0159), classical monocytes (*P* = 0.0189), and CD56dim NK cells (*P* = 0.0386), and elevations in plasma cells (*P* = 0.0112) and memory B cells (*P* = 0.019). No immune cell populations were statistically significantly elevated after FDR correction.

In AA, the most extensive shifts were observed: naïve B cells (P = 4.93 × 10□□), B cells (P = 7.75 × 10□□), CD56dim NK cells (P = 0.0252), intermediate monocytes I (P = 8.99 × 10□) and II (P = 0.0204) were all significantly decreased in AD, while plasma cells (P = 3.36 × 10), memory B cells (P = 4.58 × 10□³), CD56bright NK cells (P = 0.0227), memory CD8□ T cells (P = 0.0262), M0 macrophages (P = 1.61 × 10□³), non-classical monocytes (P = 0.0238), and neutrophils (P = 5.64 × 10□□) were elevated. Double-negative T cells (P = 0.0418, decreased) and M2 macrophages (P = 0.0396, elevated) were also nominally significant (**Supplemental Table 2**).

In CariHISP, AD was characterized by reductions in CD56dim NK cells (P = 0.0295), activated NK cells (P = 6.05 × 10□□), naïve CD4□ T cells (LM22) (P = 0.0154), naïve-to-stimulated CD4□ T cells (P = 4.76 × 10□³), and regulatory T cells (LM22) (P = 0.0124), with concurrent elevations in non-classical monocytes (P = 0.0129) and intermediate monocytes II (P = 2.81 × 10□³). Additional nominal changes reported in Supplemental Table 2 but not shown in Figure 3 were reductions in double-negative T cells (P = 0.0321), memory/naïve CD8□ T cells (P = 0.0233), and hematopoietic stem cells (P = 0.0451), and elevations in resting mast cells (P = 0.0269) and resting memory CD4□ T cells (P = 0.0183).

In PERU, almost as many significant associations as CariHISP (11 vs 12) were detected, despite having the smaller sample size: neutrophils (P = 3.63 × 10□³), non-classical monocytes (P = 9.13 × 10□□), and classical monocytes (P = 5.31 × 10□³) were elevated in AD, while activated NK cells (P = 0.0109), memory CD8□ T cells (P = 0.0118), myeloid DCs (P = 8.82 × 10□³), and plasmacytoid DCs (P = 3.19 × 10□□) were decreased. Monocytes (P = 0.0326), naïve CD4□ T cells (1M) (P = 0.0237), follicular helper T cells (P = 0.027), and CD8□ T cells (P = 0.0495) were also nominally decreased (**Supplemental Table 2**).

**Supplemental Table 2** lists the full *propeller* results for **Figure 3**.

Together, these results reveal that while peripheral immune remodeling is a consistent feature of AD, the specific cell types affected and the direction of changes vary by ancestry. AA and CariHISP shared reduced NK dim cells and elevated non-classical monocytes but otherwise differed: AA showed lymphoid depletion alongside macrophage and neutrophil expansion, whereas CariHISP showed NK and T cell depletion alongside expansion of mast cells and intermediate monocytes II. NHW showed a narrower profile of monocyte, naïve B cell, and NK dim cell reductions. AA and CariHISP exhibited the broadest alterations, spanning multiple lymphoid, myeloid, and NK cell populations. PERU showed fewer significant findings than AA and CariHISP but directionally concordant trends. These results highlight the importance of ancestry-informed analyses to better understand the immunopathology of AD and guide the development of population-sensitive biomarkers and therapeutic strategies.

### Interaction Model of Ancestry and AD Status Reveals Population-Specific Peripheral Immune Alterations in AD

The within-group analyses above show where AD-associated shifts reached significance in each group, but they do not by themselves establish that these effects differ between groups. We therefore tested interaction contrasts comparing the AD-versus-control difference in NHW with that in each of the other three groups (**Figure 4**). For each cell type, the interaction contrast was defined as the AD-versus-control difference in NHW minus the AD-versus-control difference in AA, CariHISP, or PERU, yielding three contrasts. Because all three contrasts use NHW as the reference, they test whether the AD effect in AA, CariHISP, or PERU differs from that in NHW and do not compare the other groups with one another.

Interaction effects were concentrated in CariHISP and PERU. Six cell types differed between CariHISP and NHW at FDR < 0.05, two differed between PERU and NHW, and none differed between AA and NHW (minimum FDR = 0.40). Nominal interactions (P < 0.05) were observed for 12 of 44 named cell types in CariHISP, 10 in PERU, and 4 in AA.

In CariHISP, the strongest interactions were observed for intermediate monocytes I (P = 5.25 × 10□□, FDR = 0.0118) and II (P = 8.61 × 10□□, FDR = 0.0129), activated NK cells (P = 1.40 × 10□³, FDR = 0.0138), resting memory CD4□ T cells (P = 1.53 × 10□³, FDR = 0.0138), naïve-to-stimulated CD4□ T cells (P = 2.35 × 10□³, FDR = 0.0176), and hematopoietic stem cells (P = 5.61 × 10□³, FDR = 0.0361). The reduction in intermediate monocytes I observed in NHW (P = 2.6 × 10□□) was absent in CariHISP (P = 0.675), whereas intermediate monocytes II and resting memory CD4□ T cells were elevated in CariHISP (P = 2.81 × 10□³ and P = 0.0183, respectively) and trended lower in NHW (P = 0.136 and P = 0.0578, respectively). Conversely, activated NK cells (P = 6.05 × 10□□), naïve-to-stimulated CD4□ T cells (P = 4.76 × 10□³), and hematopoietic stem cells (P = 0.0451) were reduced in CariHISP but not significantly altered in NHW (P = 0.387, 0.206, and 0.0865, respectively). Classical monocytes (P = 0.0159, FDR = 0.0897) and plasma cells (P = 0.0182, FDR = 0.0911) showed nominal interactions that did not reach FDR significance.

In PERU, classical monocytes showed the strongest interaction (P = 2.12 × 10□□, FDR = 9.55 × 10□³), reflecting opposite directions of effect: classical monocytes were reduced in NHW (P = 0.0189) but elevated in PERU (P = 5.31 × 10□³). Memory CD8□ T cells also differed (P = 1.76 × 10□³, FDR = 0.0397), with a reduction in PERU (P = 0.0118) and a nonsignificant elevation in NHW (P = 0.0849). Naïve CD4□ T cells (1M) (P = 4.50 × 10□³), plasmacytoid DCs (P = 5.20 × 10□³), and myeloid DCs (P = 5.74 × 10□³) approached FDR significance (FDR = 0.0516 for each), as did activated NK cells (P = 7.44 × 10□³, FDR = 0.0558). Each of these four was reduced in PERU (P = 0.0237, 3.19 × 10□□, 8.82 × 10□³, and 0.0109, respectively) without a corresponding change in NHW.

In contrast, no cell type showed an AD effect in AA that differed from NHW at FDR < 0.05, and all cell types shown in Figure 4 had FDR ≥ 0.67 in AA. The smallest nominal interactions in AA were for CD56bright NK cells (P = 9.90 × 10□³), M0 macrophages (P = 0.0183), naïve B cells (P = 0.0304), and B cells (P = 0.0358; FDR = 0.40 for each). Several of the cell types that were significant within AA were altered in the same direction in NHW: naïve B cells, intermediate monocytes I, and CD56dim NK cells were reduced, and plasma cells and memory B cells were elevated, in both groups. The strongest interaction overall was for the unannotated “unknown” population of the 1M reference in CariHISP (P = 5.91 × 10□□, FDR = 2.7 × 10□□), which represented less than 0.2% of the estimated composition on average and is omitted from Figure 4.

**Supplemental Table 3** lists the full interaction results for **Figure 4**.

Together, these results indicate that AD-associated immune shifts in NHW and AA differ from those in CariHISP and, to a lesser extent, in PERU. The CariHISP interactions involved NK, CD4□ T cell, intermediate monocyte, and hematopoietic stem cell compartments, whereas the PERU interactions involved classical monocytes, memory CD8□ T cells, and dendritic cells. Because these contrasts used NHW as the sole reference, they do not establish differences among AA, CariHISP, and PERU.

### Cell-Type Specific Disease-Associated Pathways Differ in Multi-Ancestry Cohorts

Building on these findings, we explored whether disease-associated transcriptional programs differ across groups by performing cell-specific differential expression (AD vs. control) within each ethnic group and summarizing the results using Hallmark pathway enrichment (**Figure 5**). The heatmaps reveal both shared and group-specific pathway signatures across immune compartments.

**Figure 5.**
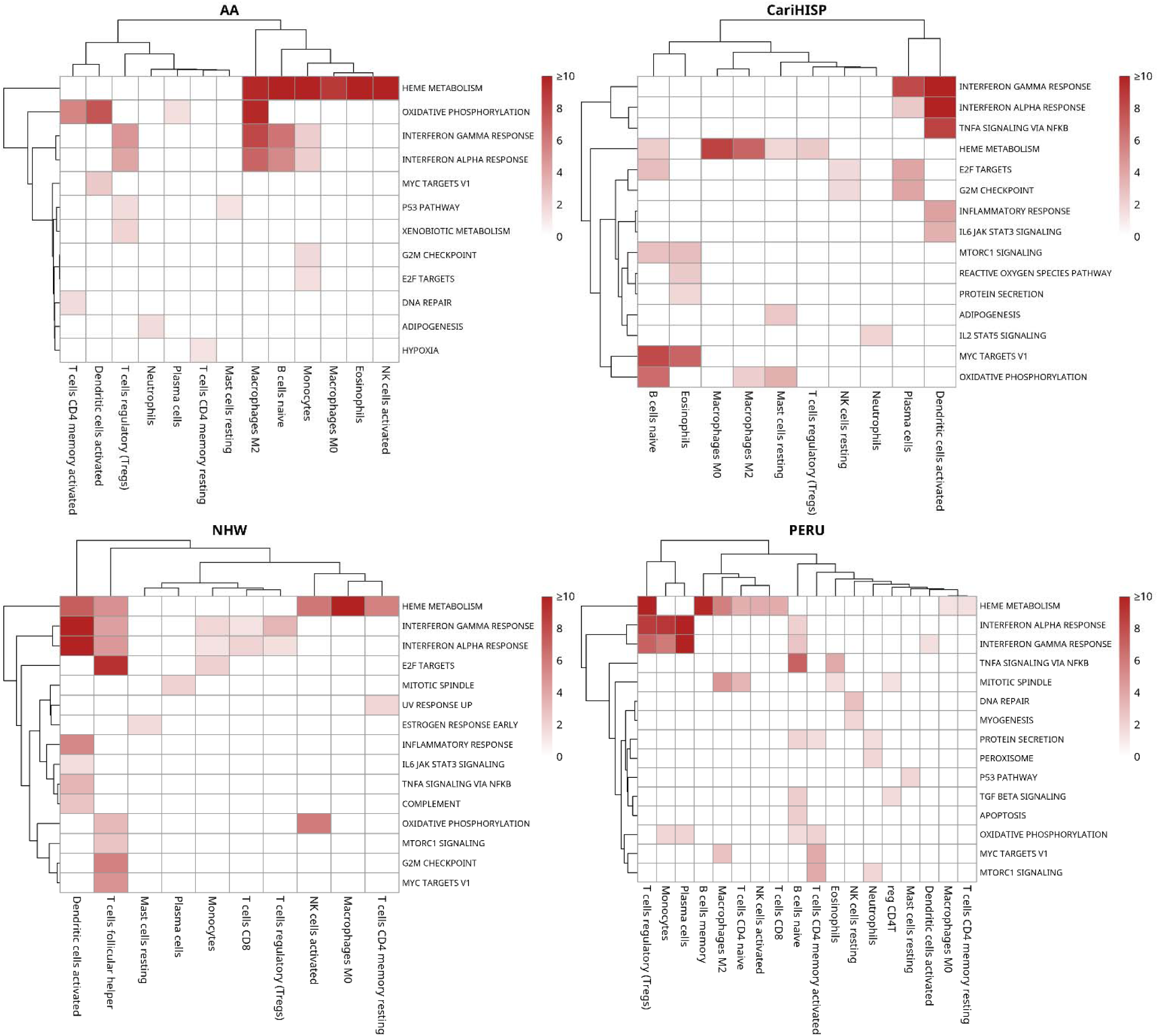
Top Enriched Pathways From Cell-Specific Differential Expression Analysis Across AD Status. Heatmaps display the top enriched Hallmark pathways (MSigDB) from cell-type-specific differential gene expression analyses comparing Alzheimer’s disease (AD) cases to cognitively normal controls, stratified by ancestry: stratified by ancestry: African American (AA, top left), Caribbean Hispanic (CariHISP, top right), non-Hispanic White (NHW, bottom left), and Peruvian (PERU, bottom right) Rows represent pathways, and columns represent immune cell types imputed from whole blood RNA-seq using CIBERSORTx. Color intensity indicates fold enrichment significance of each pathway within the respective cell type. Hierarchical clustering was applied to both rows and columns within each ancestry.

Across all groups, we observed recurrent enrichment of heme metabolism and type I/II interferon responses across multiple cell types, consistent with broad immune activation in AD. Oxidative phosphorylation was enriched in myeloid and select lymphoid populations across most groups, suggesting mitochondrial metabolic remodeling as a conserved disease feature.

In NHW, interferon responses were the dominant feature, with interferon alpha and gamma showing the strongest enrichment of any group in activated dendritic cells (−log□□P = 41.8 and 50.5, respectively), with additional signals in T follicular helper cells (4.6 and 4.3, respectively), regulatory T cells, monocytes, and CD8 T cells. Heme metabolism was strongly enriched in macrophages M0 (18.8), activated dendritic cells (7.2), activated NK cells (6.1), resting CD4□ memory T cells (5.7), and T follicular helper cells (5.1). Oxidative phosphorylation was enriched in activated NK cells (5.9) and T follicular helper cells (3.1), while E2F targets (9.3), G2M checkpoint (5.7), MYC targets V1 (4.8), and mTORC1 signaling (2.6) were enriched in T follicular helper cells. A distinct cluster of inflammatory response (5.4), complement, TNFα signaling via NF-κB, and IL6–JAK–STAT3 signaling was detected in activated dendritic cells.

Resting CD4□ memory T cells showed enrichment of UV response up (1.8), while mitotic spindle and estrogen response early appeared in plasma cells (2.0) and resting mast cells (1.7), respectively.

In AA, heme metabolism was the dominant signal, most strongly enriched in eosinophils (−log□□ P = 24.0), activated NK cells (13.1), and naïve B cells (12.6), with additional enrichment in monocytes, macrophages M2, and macrophages M0. Oxidative phosphorylation was prominent in macrophages M2 (9.6), activated dendritic cells (7.7), and activated CD4 memory T cells (5.5). Interferon gamma and alpha responses were concentrated in macrophages M2, naïve B cells, monocytes, and regulatory T cells. Additional signatures included adipogenesis in neutrophils, hypoxia in resting CD4 memory T cells, E2F targets and G2M checkpoint in monocytes, MYC targets V1 in activated dendritic cells, p53 pathway in regulatory T cells and resting mast cells, xenobiotic metabolism in regulatory T cells, and DNA repair in activated CD4 memory T cells—collectively suggesting metabolic stress and cell-cycle dysregulation across myeloid and select lymphoid populations.

In CariHISP, heme metabolism was strongly enriched in macrophages M0 (8.6) and macrophages M2 (7.0), with weaker signals in regulatory T cells (2.5), naïve B cells (2.3), and resting mast cells (1.8). MYC targets V1 was strongly enriched in naïve B cells (8.1) and eosinophils (6.9). Oxidative phosphorylation was enriched across three cell types: naïve B cells (6.6), resting mast cells (3.5), and macrophages M2 (2.2). Interferon alpha and gamma responses were concentrated in activated dendritic cells (18.6 for both) and plasma cells (2.4 and 8.2, respectively), with activated dendritic cells also showing TNFα signaling via NF-κB (8.6), inflammatory response (4.2), and IL6–JAK–STAT3 signaling (3.6). Additional CariHISP-specific programs included E2F targets in plasma cells (4.1), naïve B cells, and resting NK cells, G2M checkpoint in plasma cells and resting NK cells, mTORC1 signaling in eosinophils and naïve B cells, reactive oxygen species and protein secretion in eosinophils, adipogenesis in resting mast cells, and IL2–STAT5 signaling in neutrophils—indicative of combined proliferative and proteostasis stress.

In PERU, the pathway profile was the broadest in terms of cell type coverage, with enriched pathways detected in eighteen cell types. Interferon alpha and gamma responses were enriched across four and five cell types, respectively, with the strongest signals in plasma cells (12.3 and 15.6), monocytes (9.0 and 6.0), and regulatory T cells (8.8 and 7.0). Heme metabolism spanned eight cell types, most prominently regulatory T cells (21.5), memory B cells (9.7), and macrophages M2 (5.7). MYC targets V1 was enriched in activated CD4 memory T cells (3.8) and macrophages M2 (2.8). DNA repair was notably enriched in resting NK cells (2.8). TNFα signaling via NF-κB was detected in naïve B cells (7.0) and eosinophils (3.9), and mTORC1 signaling in activated CD4 memory T cells (3.9) and neutrophils (1.8). Additional PERU-specific signals included mitotic spindle in macrophages M2 (4.8), naïve CD4 T cells, eosinophils, and regulatory CD4 T cells, TGF-β signaling in naïve B cells and regulatory CD4 T cells, apoptosis in naïve B cells, protein secretion in naïve B cells, neutrophils, and activated CD4 memory T cells, myogenesis in resting NK cells, peroxisome in neutrophils, and p53 pathway in resting mast cells—reflecting the most widespread array of immune-metabolic perturbations among the four groups despite the smaller sample size.

**Supplemental Table 4** shows the pathway enrichment results of **Figure 5**. **Supplemental Figures 1-6** show pathway enrichment results in the KEGG pathway database, WikiPathways, C2 Canonical Pathways, C3 microRNA targets, C5 Human Phenotype Ontology, and C7 immunologic signature gene sets respectively. **Supplemental Tables 5-10** show the enrichment analyses results of each of **Supplemental Figures 1-6** respectively.

Together, these analyses indicate that AD engages a common set of immune-metabolic programs—centered on heme metabolism, interferon signaling, and oxidative phosphorylation—across cell types and ancestries, but the strength, breadth, and distribution of these programs vary considerably by group. The convergence on heme, interferon, and mitochondrial pathways supports a model of systemic immune activation with metabolic stress, while the group-specific profiles—particularly the broad cell type coverage in PERU and the dendritic cell–centered signatures in NHW and CariHISP—underscore the importance of population-informed analyses to elucidate mechanistic diversity and guide targeted biomarker and therapeutic development.

## Discussion

Our study provides a comprehensive view of peripheral immune changes associated with Alzheimer’s disease (AD) across four ethnic groups—non-Hispanic White (NHW), African American (AA), Caribbean Hispanic (CariHISP), and Peruvian (PERU)—and highlights both shared and group-specific features of immune remodeling. Using high-resolution deconvolution of whole-blood RNA-seq (CIBERSORTx/1M-scBloodNL), composition-aware modeling (*propeller*), and cell-type-specific differential expression with pathway enrichment, we delineate differences in baseline immune setpoints as well as disease-associated shifts that vary by group.

At baseline, immune cell composition differed significantly across groups after adjusting for age and sex, with three broad patterns emerging (**Figure 1**). Th2 CD4□ T cells and cytotoxic CD4□ T cells were highest in NHW with intermediate levels in CariHISP and AA and lower levels in PERU. Conversely, plasma cells and non-classical monocytes were most abundant in CariHISP and PERU with relative depletion in NHW and AA. A third set of cell types—regulatory CD4□ T cells, plasmacytoid DCs, intermediate monocyte I, megakaryocytes, and double-negative T cells—showed broadly similar distributions across NHW, AA, and CariHISP but marked divergence in PERU. These patterns indicate that ancestry captures meaningful variation in peripheral immune architecture that could influence how AD manifests in blood.

When evaluating AD versus control across all groups combined, we observed a striking pattern of lymphoid and intermediate monocyte depletion alongside neutrophil and non-classical monocyte expansion (**Figure 2**). Eight cell types reached FDR significance, with the strongest associations for naïve B cells, non-classical monocytes, and neutrophils: naïve B cells decreased and non-classical monocytes and neutrophils increased in AD. Neutrophils, non-classical monocytes, and plasma cells were the only cell types significantly elevated. Additional reductions were observed in CD56dim NK cells, B cells, intermediate monocyte I, and double-negative T cells. This pattern of concurrent lymphoid contraction and neutrophil expansion is consistent with chronic inflammatory remodeling in AD,^41^ potentially reflecting exhaustion or consumption of adaptive immune populations alongside neutrophil mobilization driven by circulating inflammatory mediators and Aβ-related immune stimulation.^42,43^

Critically, within-group analyses showed that the magnitude, breadth, and direction of AD-associated changes differ across groups (**Figure 3**), and interaction tests supported group-specific effects in CariHISP and PERU relative to NHW (**Figure 4**).. In NHW, AD was associated with reductions in four cell types—naïve B cells, classical monocytes, CD56dim NK cells, and intermediate monocytes I—with only plasma cells and memory B cells nominally elevated, suggesting a pattern of monocyte-centered depletion. In AA, the most extensive shifts were observed: naïve B cells, B cells, NK dim cells, and intermediate monocytes I and II were decreased (as were double-negative T cells, nominally), while plasma cells, NK bright cells, memory CD8□ T cells, macrophages M0, memory B cells, non-classical monocytes, and neutrophils were elevated (as were M2 macrophages, nominally), indicating a complex remodeling in which lymphoid depletion coexists with macrophage expansion and neutrophil mobilization. In CariHISP, AD showed a more mixed profile: CD56dim NK cells, activated NK cells, naïve CD4□ T cells, naïve-to-stimulated CD4□ T cells, and regulatory T cells were reduced, along with double-negative T cells, memory/naïve CD8 T cells, and hematopoietic stem cells at nominal significance, while non-classical monocytes and intermediate monocytes II were elevated, along with resting mast cells and resting memory CD4□ T cells at nominal significance, suggesting NK and T cell depletion coupled with myeloid and innate immune expansion. In PERU, fewer significant changes were detected than in AA and CariHISP—neutrophils, non-classical monocytes, and classical monocytes elevated; activated NK cells, memory CD8□ T cells, myeloid DCs, and plasmacytoid DCs decreased, along with monocytes, naïve CD4□ T cells, follicular helper T cells, and CD8□ T cells at nominal significance—consistent with limited statistical power but directionally concordant trends. These results underscore the importance of interaction-aware modeling; simple pooling would dilute or mask population-specific biology.

Interaction analyses of AD status by ethnic group refined the picture provided by the within-group analyses (**Figure 4**). Differences from NHW were supported in CariHISP, where 6 cell types reached FDR significance, and to a lesser degree in PERU (2 with FDR significance and 4 were close). No cell type differed between AA and NHW, even though AA showed the largest number of within-group associations (**Figure 3**). Because the AA and NHW cohorts were similar in size, this suggests that many AA associations reflect effects of similar direction and magnitude to those in NHW, including reductions in naïve B cells, intermediate monocytes I, and NK dim cells, that crossed the significance threshold in one group but not the other. Interaction tests have less power than within-group tests, so modest AA-specific differences cannot be excluded. Still, a cell type being significant in one group and not another is not itself evidence of heterogeneity, and group-specific interpretations are best supported where the interaction tests agree.

CariHISP interactions point to differences in NK, CD4□ T cell, and intermediate monocyte compartments. Intermediate monocytes I were reduced in NHW but unchanged in CariHISP, whereas intermediate monocytes II and resting memory CD4□ T cells were elevated in CariHISP and trended lower in NHW. Activated NK cells, naïve-to-stimulated CD4□ T cells, and hematopoietic stem cells were reduced in CariHISP only, consistent with the NK and T cell depletion coupled with myeloid expansion described above. The near-absence of intermediate monocytes I at baseline in CariHISP (**Figure 1**) may limit the ability to detect a further decrease, suggesting that baseline immune setpoints could partly shape the apparent AD effect.^31,32^ The opposing behavior of the two intermediate monocyte subsets is also consistent with reports that monocyte activation in early AD is heterogeneous,^25,42^ and suggests that monocyte remodeling in AD may be both subset– and population-dependent.

In PERU, the strongest interaction involved classical monocytes, which were reduced in NHW and elevated in PERU. This sign difference means that a classical monocyte signal derived in one group would point in the opposite direction in the other. The remaining PERU interactions, reflecting lower memory CD8 T cells, dendritic cells, and activated NK cells, had FDR values of 0.040 to 0.056 and should be considered suggestive given the small PERU cohort (n = 88) and its younger age distribution (**Table 1**). Together, these results strengthen the case for population-specific thresholds or stratification in blood-based immune biomarkers, particularly for monocyte– and NK-derived signals.

**Table 1.** presents the baseline demographic and genetic characteristics of participants across the non-Hispanic White (NHW, n = 240), African American (AA, n = 234), Caribbean Hispanic (CariHISP, n = 294), and Peruvian (PERU, n = 88) cohorts. Sex distribution differed significantly across cohorts (*P* < 0.001), with a higher proportion of female participants in the AA and CariHISP groups compared to NHW and PERU. Mean age at exam and age of onset also differed significantly (*P* < 0.001), with PERU participants being slightly younger on average. APOE genotype frequencies varied substantially across cohorts (*P* < 0.001), with the ε3/ε4 and ε4/ε4 genotypes more frequent among AA participants compared to other groups. 1 NHW individual did not have an APOE genotype. The proportion of Alzheimer’s disease (AD) cases and controls was well balanced across all cohorts (*P* = 0.938), indicating good case–control matching for downstream analyses.

|  |  | *NHW* | *AA* | *CariHISP* | *PERU* | *p* |
|---|---|---|---|---|---|---|
| ***Sample Size*** |  | 240 | 234 | 294 | 88 |  |
| ***Sex*** | **Female** | 129 (53.8) | 169 (72.2) | 210 (71.4) | 51 (58.0) | <0.001 |
|  | **Male** | 111 (46.2) | 65 (27.8) | 84 (28.6) | 37 (42.0) |  |
| ***Age of Exam*** |  | 77.59 (7.30) | 76.54 (7.80) | 77.91 (7.59) | 72.88 (7.95) | <0.001 |
| ***Age of Onset*** |  | 74.31 (6.83) | 75.74 (6.36) | 75.21 (7.50) | 69.20 (9.16) | <0.001 |
| ***APOE Genotype*** | **2\|2** | 2 (0.8) | 3 (1.3) | 0 (0.0) | 0 (0.0) | <0.001 |
|  | **2\|3** | 19 (7.9) | 37 (15.8) | 30 (10.2) | 4 (4.5) |  |
|  | **2\|4** | 4 (1.7) | 16 (6.8) | 9 (3.1) | 0 (0.0) |  |
|  | **3\|3** | 132 (55.2) | 63 (26.9) | 173 (58.8) | 67 (76.1) |  |
|  | **3\|4** | 71 (29.7) | 87 (37.2) | 66 (22.4) | 15 (17.0) |  |
|  | **4\|4** | 11 (4.6) | 28 (12.0) | 16 (5.4) | 2 (2.3) |  |
|  | **Missing** | 1 (0.4) | 0 (0.0) | 0 (0.0) | 0 (0.0) |  |
| ***Disease Status*** | **Control** | 119 (49.6) | 119 (50.9) | 151 (51.4) | 47 (53.4) | 0.938 |
|  | **AD** | 121 (50.4) | 115 (49.1) | 143 (48.6) | 41 (46.6) |  |

Several caveats apply to these interaction analyses. First, all contrasts used NHW as the reference, so differences among AA, CariHISP, and PERU were not formally tested, and no global test of heterogeneity was performed. Second, ancestry groups correspond to recruitment cohorts and CIBERSORTx was run separately within each, so interactions may also capture site, batch, age, sex, or environmental differences that cannot be separated from ancestry in this design. The strongest interaction overall was for a small unannotated cell population in the reference, which hints that deconvolution may behave differently across groups.

Cell-type-specific differential expression with Hallmark pathway enrichment further revealed convergent programs and divergent emphases across groups (**Figure 5**). Across all groups, heme metabolism, type I/II interferon responses, and oxidative phosphorylation were recurrently enriched, consistent with immune activation coupled to mitochondrial and redox remodeling in AD. In NHW, interferon alpha and gamma responses dominated, with the strongest enrichment of any group concentrated in activated dendritic cells and more modest signals in T follicular helper cells and regulatory T cells, alongside strong heme metabolism signals in macrophages M0 and activated dendritic cells. In AA, heme metabolism was the most prominent signature, with particularly strong enrichment in eosinophils, activated NK cells, and naïve B cells, accompanied by oxidative phosphorylation in macrophages M2, activated dendritic cells, and activated CD4□ memory T cells—aligning with the compositional myeloid and macrophage shifts observed. CariHISP showed the strongest MYC targets V1 enrichment, concentrated in naïve B cells and eosinophils, with oxidative phosphorylation in naïve B cells, resting mast cells, and macrophages M2; additional programs including E2F targets, G2M checkpoint, mTORC1 signaling, reactive oxygen species, and IL2–STAT5 signaling were enriched in lymphoid, granulocyte, and mast cell populations, and IL6–JAK–STAT3 signaling in activated dendritic cells, suggestive of combined proliferative and proteostasis stress. PERU displayed the broadest pathway landscape in terms of cell type coverage: heme metabolism spanned eight cell types, interferon alpha and gamma responses spanned four and five cell types, respectively, and additional signals included mitotic spindle, TNFα signaling via NF-κB, TGF-β signaling, apoptosis, and DNA repair across lymphoid, myeloid, and NK populations—reflecting the most widespread array of immune-metabolic perturbations despite the smaller sample size. These pathway-level results reinforce that AD engages a shared immune-metabolic axis while the distribution and intensity of responses vary by group.

Our findings are consistent with reports of systemic inflammation and immune dysregulation in AD,^44,45^ but they refine this view by demonstrating population-dependent manifestations. The observation that AA and CariHISP show the broadest compositional changes, with reduced NK dim cells and elevated non-classical monocytes in both, while NHW displays a narrower profile of monocyte, naïve B cell, and CD56dim NK cell depletion, and PERU shows fewer significant cell-type shifts than AA and CariHISP despite the broadest cell type coverage in pathway enrichment, echoes known differences in immune architecture across populations,^31^ and suggests that baseline immune setpoints may shape disease-associated responses.^32,46^ This has direct implications for biomarker development and therapeutic design: platforms that rely on lymphocyte– or monocyte-derived signals, or that target interferon/mitochondrial pathways, may require population-specific thresholds or stratification to achieve accuracy and efficacy.

This study also advances the conceptual framework of AD as a systemic disorder. The detection of robust peripheral immune signals across multiple cell types supports a model in which AD-related pathology involves bidirectional communication between the central nervous system and peripheral immune compartments.^20^ The observed depletion of lymphoid and intermediate monocyte populations, alongside expansion of neutrophils and non-classical monocytes in the pooled analysis, suggests that peripheral immune cells may not merely reflect CNS pathology but actively participate in disease-associated immune remodeling. The lymphoid contraction may reflect chronic antigen-driven exhaustion or redistribution of immune cells to sites of neuroinflammation,^30,47^ while neutrophil expansion is consistent with reports of neutrophil hyperactivation and brain infiltration in AD.^26–28^

Despite these insights, several limitations warrant discussion. First, the deconvolution and Hi-Res transcriptomic reconstruction methods rely on reference profiles derived from European-descent individuals, which may introduce biases in estimating cell types or gene expression in non-European populations. We also note that more accurate techniques for immune cell profiling are available, but are generally cost-prohibitive for studies of this scale. While our cross-sectional design identifies associations between AD status and immune profiles, causal relationships remain speculative. Longitudinal studies are needed to determine whether peripheral immune changes precede clinical onset or result from disease pathology and/or neurodegeneration. Lastly, although our sample represents one of the largest multi-population blood transcriptomic datasets in AD, the modest group sizes within each group constrain statistical power, particularly for rare cell types and the PERU cohort.

In conclusion, our findings demonstrate that peripheral immune remodeling in AD is not monolithic but varies meaningfully across populations. AD is characterized by depletion of lymphoid and intermediate monocyte populations with concurrent neutrophil and non-classical monocyte expansion. Ancestry-stratified analyses showed this was not uniform: broad immune remodeling was most extensive in African American and Caribbean Hispanic individuals, neutrophil expansion was significant only in African American and Peruvian participants, and non-Hispanic White participants showed more limited, monocyte-centered reductions. These insights offer a foundation for developing precision immunotherapies and blood-based biomarkers that account for genetic and immunological background, ultimately improving outcomes for global populations affected by Alzheimer’s disease.

## Acknowledgements

This work made use of the High-Performance Computing Resource in the Core Facility for Advanced Research Computing at Case Western Reserve University. Data for this study were prepared, archived, and distributed by the National Institute on Aging Alzheimer’s Disease Data Storage Site (NIAGADS) at the University of Pennsylvania (U24-AG041689), funded by the National Institute on Aging. We thank the individuals who participated in the Multi-Ancestry Genomics, Epigenomics, and Transcriptomics of Alzheimer’s (MAGENTA) study, which was supported by the National Institute on Aging (NIA).

## Author Contributions

YM: Design, analysis, and manuscript preparation.

## Abbreviations

AA: African American
Aβ: amyloid-β
AD: Alzheimer’s disease
GWAS: genome-wide association study
HISP: Hispanic
IFN-γ: interferon-gamma
LPS: lipopolysaccharide
NHW: Non-Hispanic White

