## Supplemental Figures for "Peripheral Immune Cell Shifts Associated With Alzheimer’s Disease Vary in Multi-Ancestry Cohorts"

**Supplemental Figure 1**
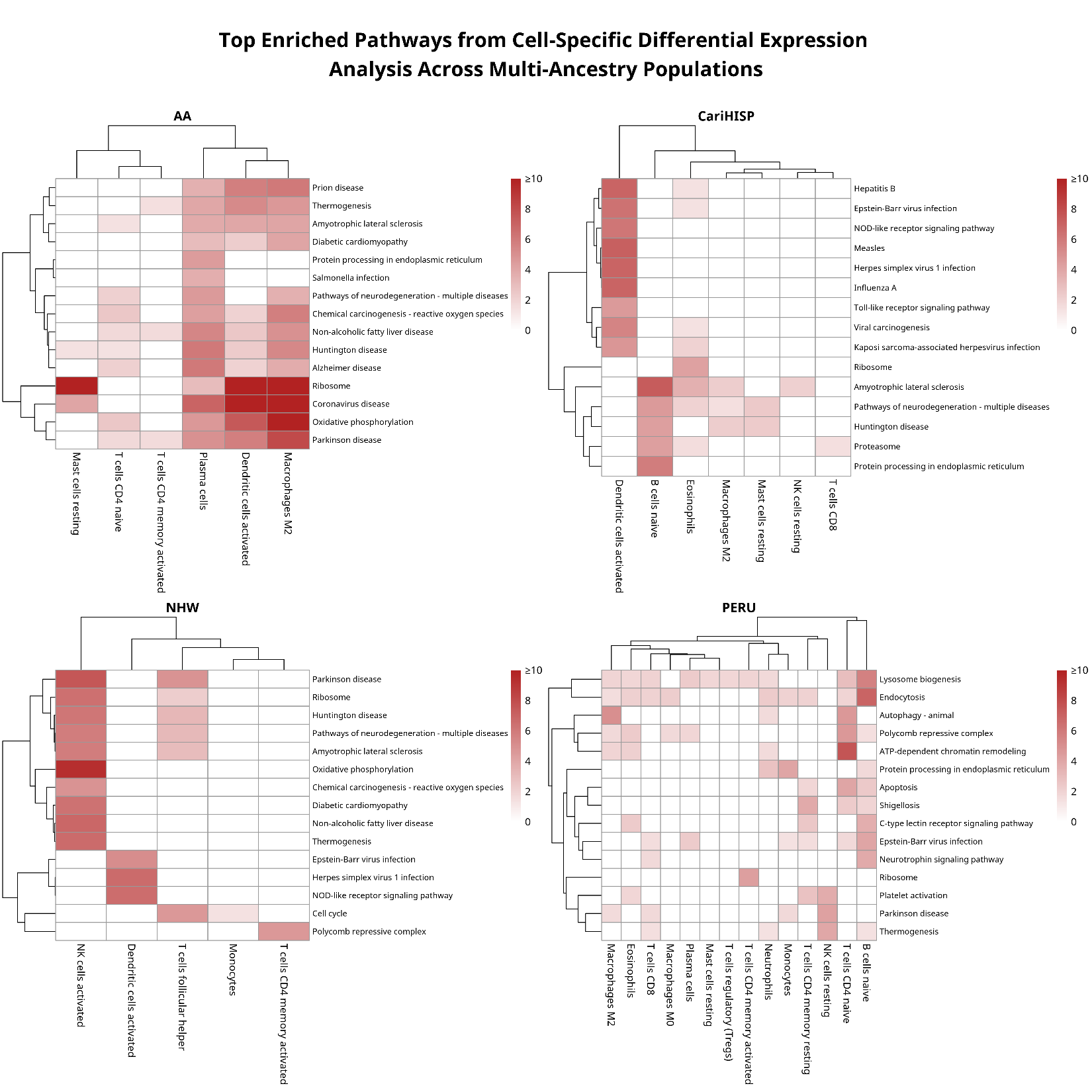

**Supplemental Figure 2**
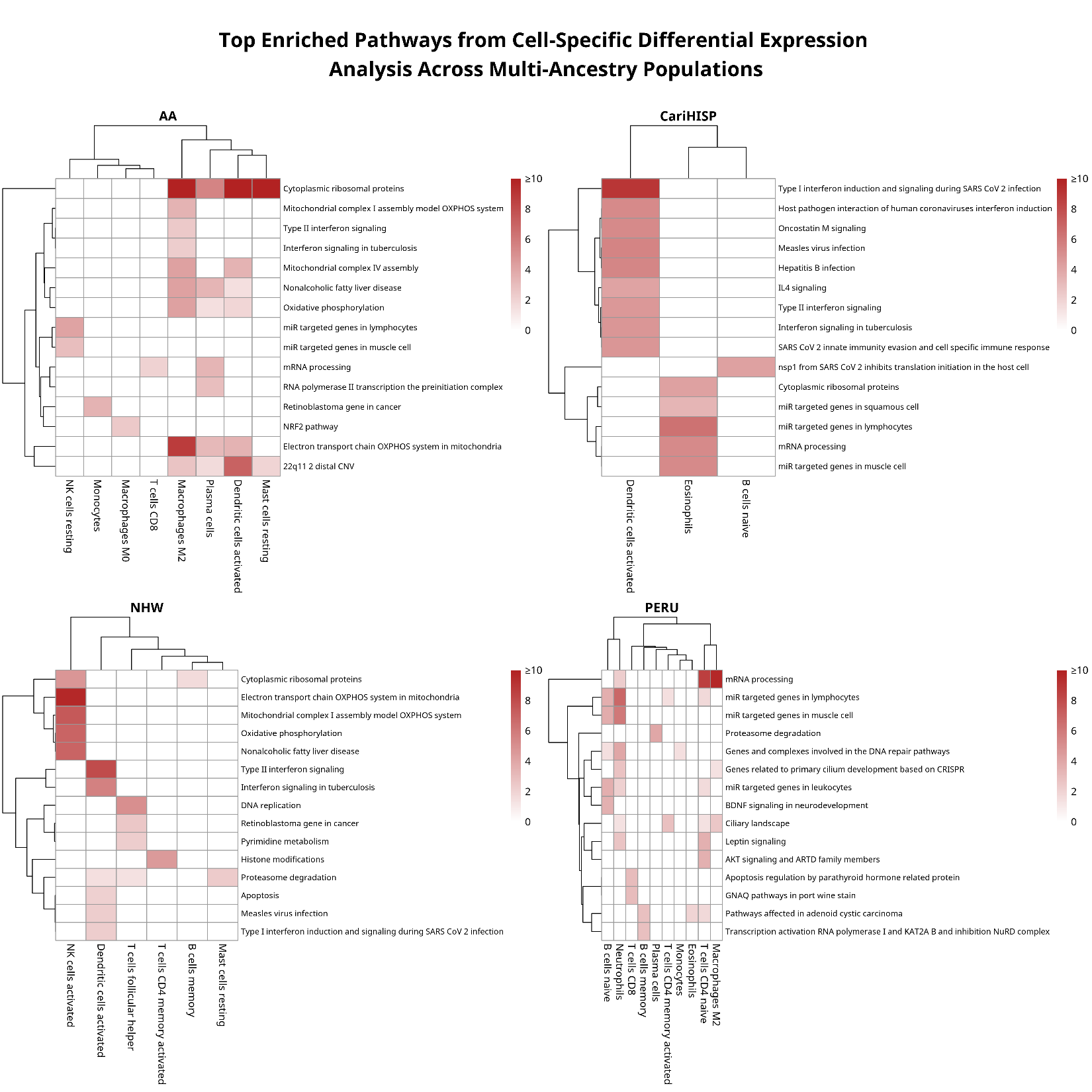

**Supplemental Figure 3**
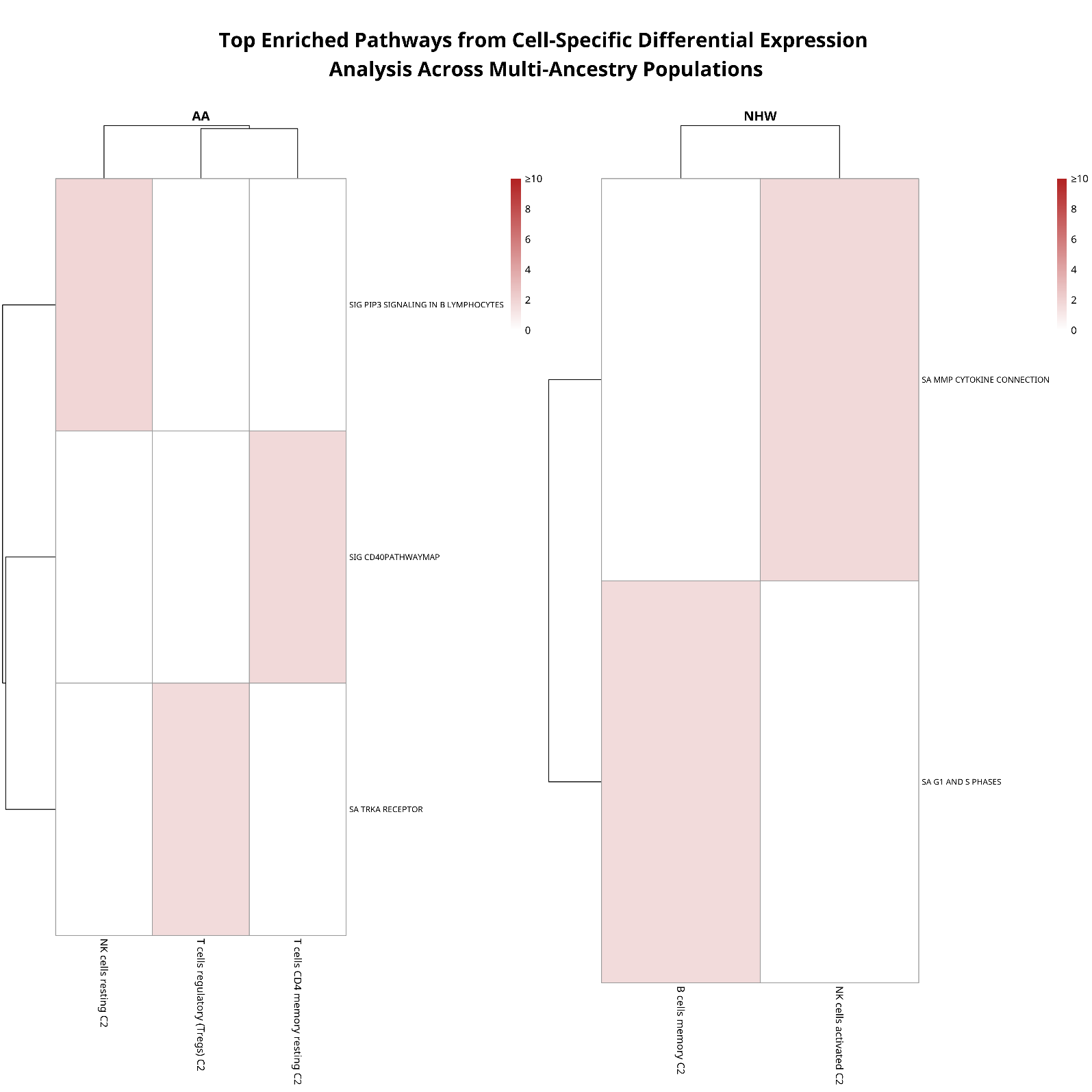

**Supplemental Figure 4**
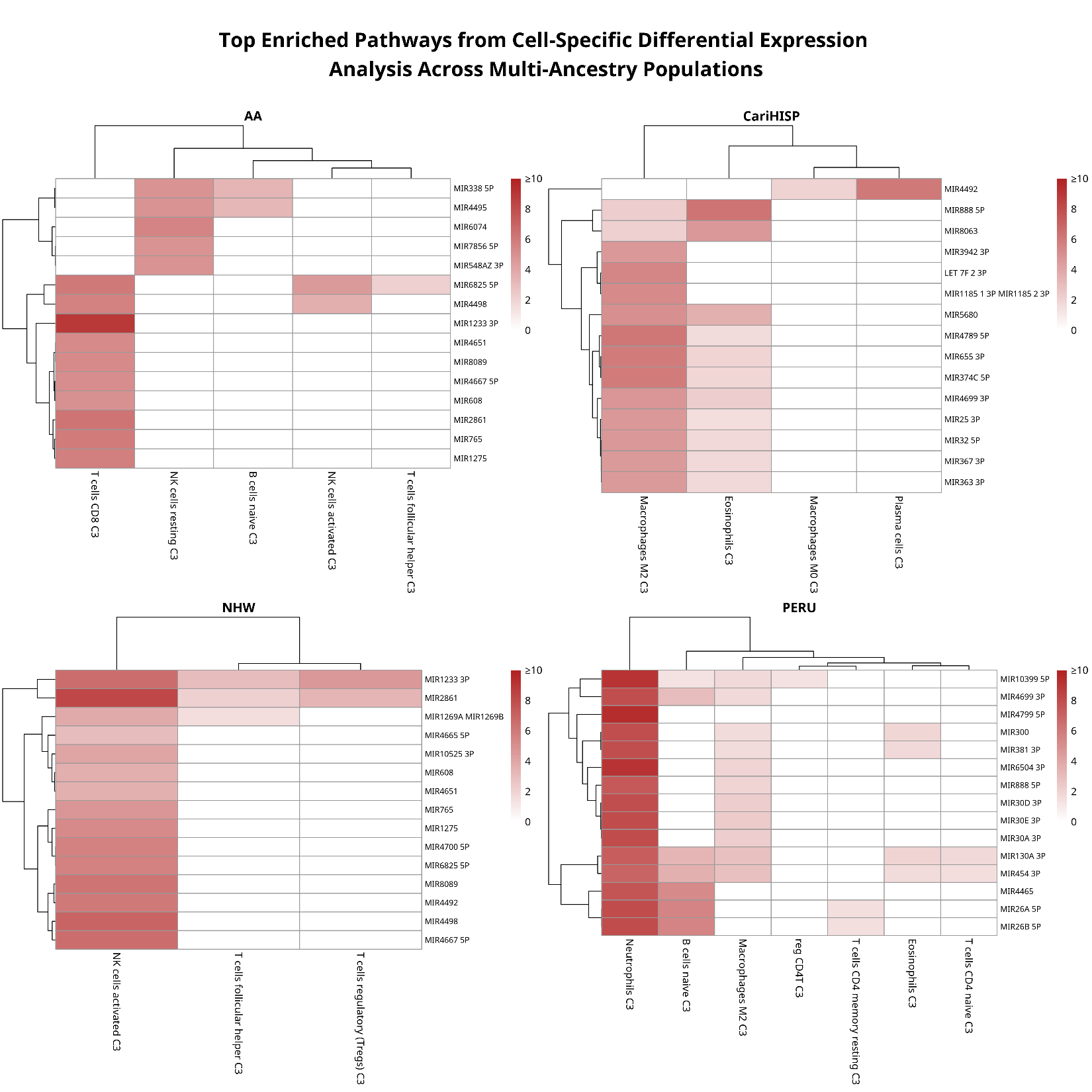

**Supplemental Figure 5**

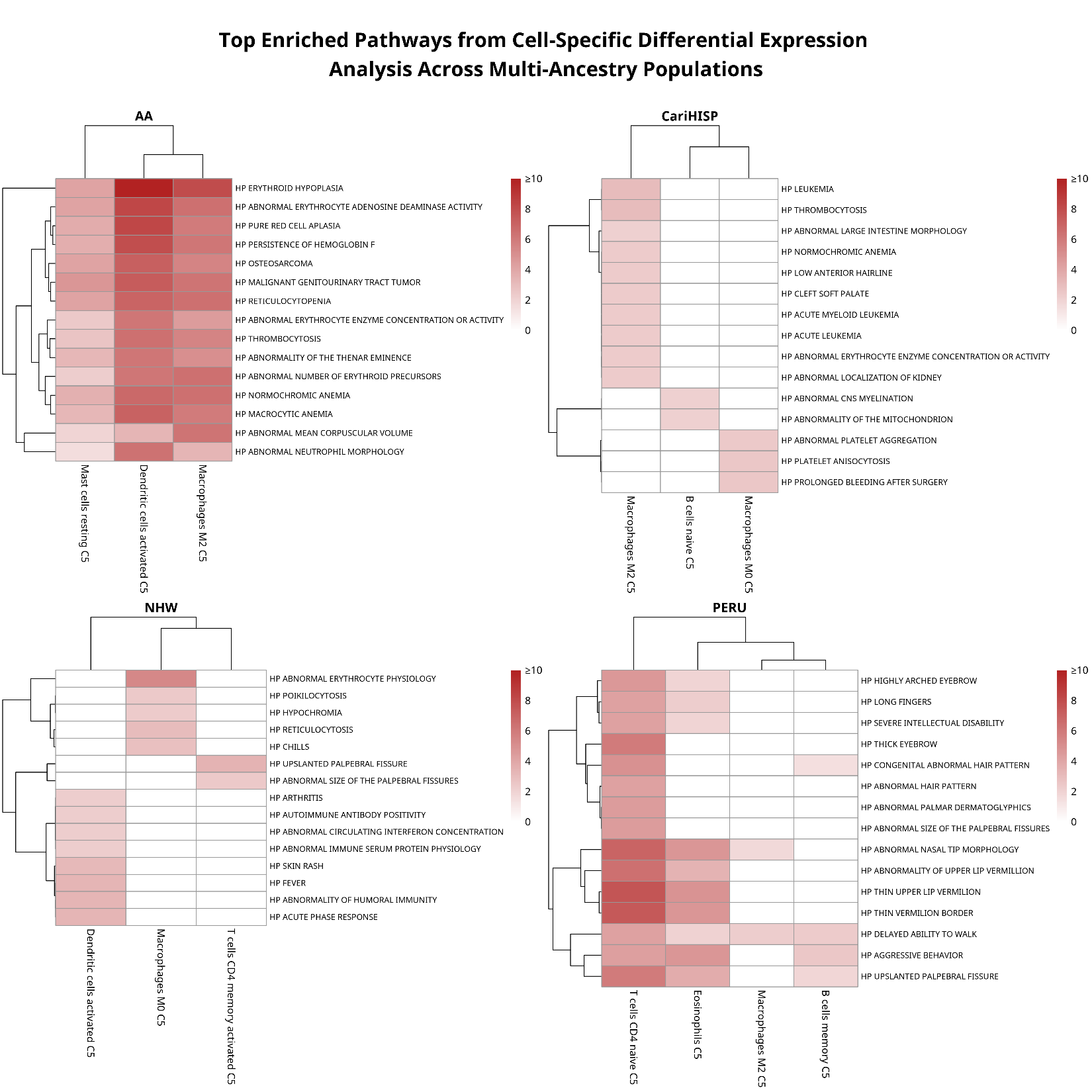

**Supplemental Figure 6**

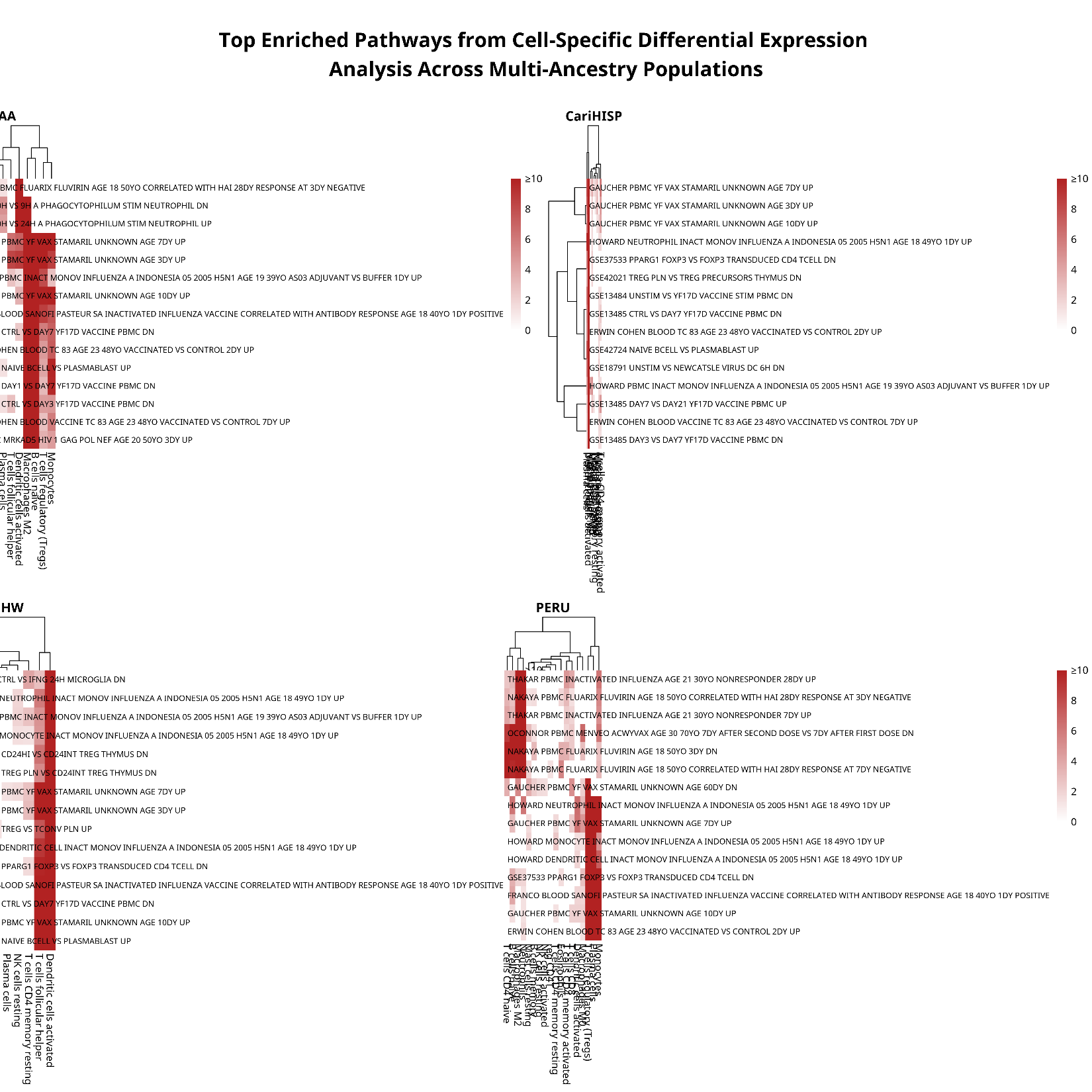
